# Losartan rewires ovarian cancer tumor-immune microenvironment and suppresses IGF-1 to amplify chemo-immunotherapy sensitivity

**DOI:** 10.1101/2023.08.31.555783

**Authors:** Yao Sun, Zhenzhen Yin, Limeng Wu, Changli Yue, Yanling Zhang, Sonu Subudhi, Pinji Lei, Alona Muzikansky, Luo Zhang, Bo R. Rueda, Rakesh K. Jain, Lei Xu

**Author notes:** Corresponding authors: Lei Xu (Edwin L. Steele Laboratories, Department of Radiation Oncology, Cox-7, Massachusetts General Hospital, Boston, Massachusetts 02114.). Y.S. and Z.Y. contributed equally to this work.

## Abstract

Ovarian cancer (OvCa) is the most lethal of the gynecologic malignancies. Immune checkpoint inhibitors, which have revolutionized the treatment of multiple malignancies, have had limited efficacy in OvCa patients. Here, using syngeneic OvCa models and genetic and pharmacologic perturbations, we discovered that losartan – a widely prescribed anti-hypertensive drug – exhibits dual effects on both the tumor microenvironment and cancer cells to sensitize OvCa to chemo-immunotherapy. Specifically, losartan treatment i) reprograms the tumor microenvironment leading to increased vascular perfusion, and thus enhances drug delivery and immune effector cell intratumoral infiltration and function; and ii) rewires the OvCa cells by suppressing the IGF-1 signaling, resulting in enhanced chemosensitivity. As a result of the combined tumor and stromal effects, losartan treatment enhances the efficacy of chemo-immunotherapy in OvCa models. The safety and low cost (less than $1-2/day) of losartan warrant rapid translation of our findings to patients with OvCa.

## Introduction

Ovarian Cancer (OvCa) is the most lethal gynecologic malignancy worldwide, with an estimated 19,710 new diagnoses and 13,270 deaths in the U.S. in 2023 ^1^. Following initial debulking surgery, OvCa patients generally receive a chemotherapy regimen that includes a platinum complex (carboplatin or cisplatin) and a taxane (paclitaxel or docetaxel) ^2^. However, despite initial responsiveness, the majority of patients with advanced OvCa eventually relapse with resistant disease ^2^. Outcomes for patients with platinum-resistant OvCa are poor, and options remain limited, making the development of new OvCa treatment a high clinical priority.

The use of immune checkpoint inhibitors (ICIs), such as antibodies against programmed cell death-1 (PD-1) and cytotoxic T-lymphocyte-associated protein 4 (CTLA-4), has revolutionized the therapy of several malignancies. Unfortunately, ICIs have had limited efficacy in OvCa, with objective response rates in only 10 to 15% of patients ^3,4^. Prevailing preclinical and clinical data indicate that abnormalities in the tumor microenvironment (TME) contribute to immunosuppression and dictate the outcome of ICIs ^5-7^. OvCa harbors an abnormal TME – rich in fibroblasts and extracellular matrix (ECM)^8,9^. The abundant ECM can generate a physical force known as”solid stress” that can compress the blood and lymphatic vessels ^10-12^. The resulting reduction in tumor blood perfusion can impair the delivery of drugs, including ICIs and immune cells, and cause hypoxia and low pH ^11,12^. Hypoxia and low pH also contribute to immunosuppression and hypoxic cells are more aggressive and more resistant to radiotherapy and chemotherapeutics that require oxygen to be effective ^13-20^. Developing a therapeutic approach that can normalize the TME is an important need to improve immunotherapy for patients with OvCa.

In our attempts to normalize the TME, we discovered that angiotensin II receptor 1 (AT1) is a key regulator of the compressive solid stress ^21,22^. The renin-angiotensin system (RAS) is known for its pivotal role in maintaining cardiovascular homeostasis as well as fluid and electrolyte balance ^23^. Angiotensin II (AngII) was initially discovered as a vasoconstrictor but is also known to contribute to the formation of ECM molecules and regulate inflammation ^23,24^. We showed in OvCa models that losartan treatment reduced matrix content and re-opened compressed vessels. As a result, blood perfusion increased, leading to enhanced drug delivery and improved chemotherapy efficacy. Furthermore, our retrospective analysis demonstrated that treatment with angiotensin blockers is associated with a 30-month overall survival benefit in OvCa patients receiving chemotherapy ^25^. As a natural extension of this study, we are now asking the question: Can losartan reprogram the immunosuppressive TME of OvCa to enhance immunotherapy alone or with chemotherapy?

Here, we report that losartan treatment, i) via normalizing the TME, improved anti-PD1 antibody delivery, and increased immune cell infiltration and function, and ii) via regulating the IGF pathway, increased OvCa cell chemosensitivity. By exerting dual effects on both the TME and cancer cells, the administration of losartan enhanced the efficacy of chemo-immunotherapy in ovarian cancer models. These findings offer new hope for improving treatment for this deadly disease.

## Results

### Losartan treatment reverses immune suppression in syngeneic OvCa models

To examine the impact of losartan on both cancer cells and TME, we performed transcriptome profiling of the SKOV3 ip1 human OvCa tumors grown i.p. in mice treated with or without losartan. We aligned bulk tumor RNA sequencing (RNASeq) data to human (GRCh37/hg19) and mouse (NCBI37/mm9) genomes, respectively. We identified 36 human genes and 109 mouse genes that were differentially expressed between control and losartan-treated tumors (FDR-adjusted p<0.05 and Log2FC>2, Supplementary Table S1). Gene Set Enrichment Analysis (GSEA) revealed that in the human tumor cells, losartan treatment leads to inhibition of pathways driving cancer progression, ECM, and fibrogenic Transforming Growth Factor (TGF)-β and Wnt signaling. In the mouse stromal compartment, losartan treatment enhances immune response pathways, such as antigen presentation, natural killer (NK) cell cytotoxic activity, chemokine signaling involved in recruiting immune cells, Janus kinases (JAK)/Signal transducer and activator of transcription protein (STAT) signaling that plays a critical role in immunity and tumor progression, and Toll-like receptor (TLR) that serves as a link between innate and adaptive immunity, respectively ^26^ (Fig 1A-B).

**Figure 1.**
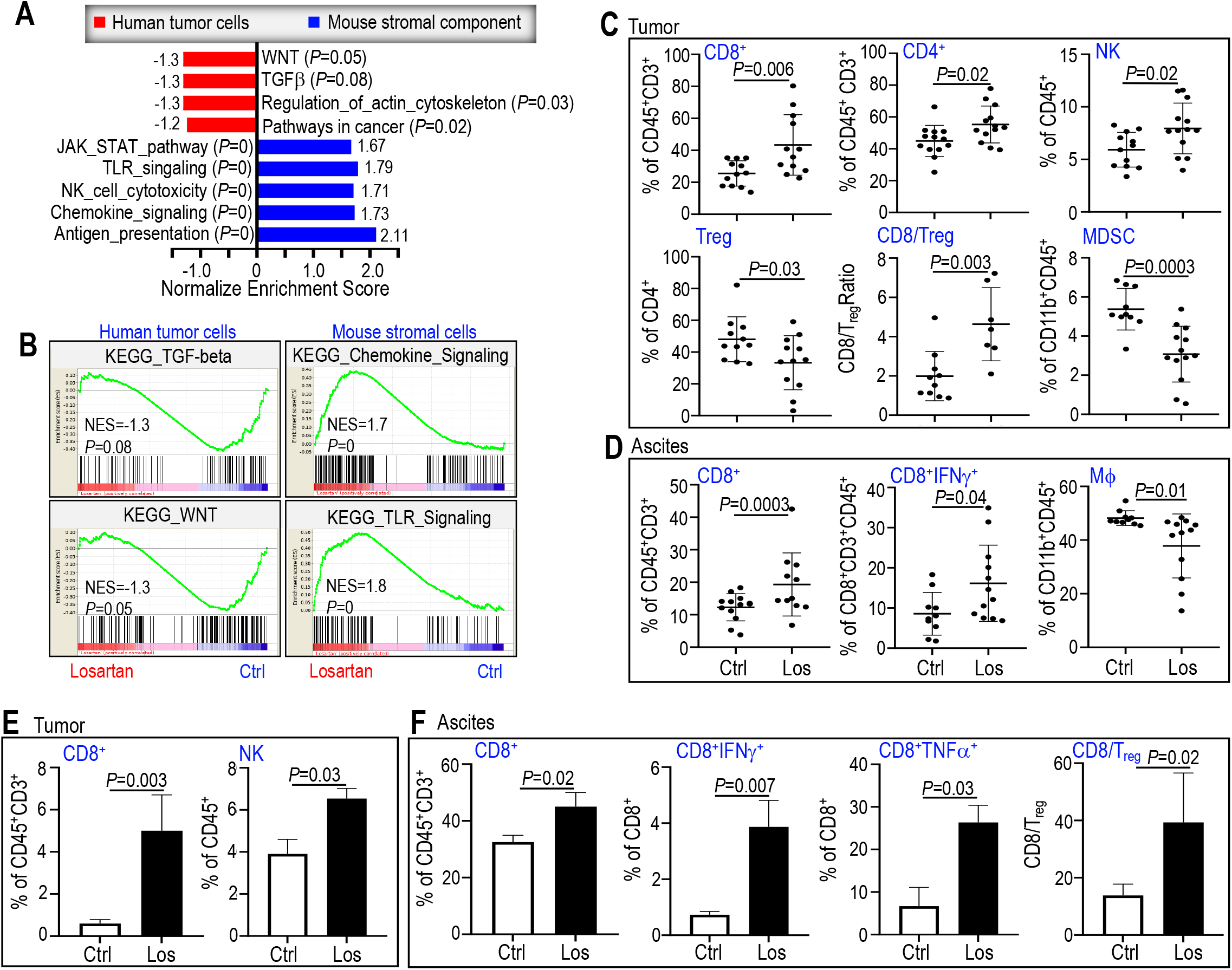
Losartan treatment reprograms the TME from immunosuppressive to immunostimulatory in OvCa models. **(A-B)** RNASeq analysis of human SKOV3ip1 OvCa xenograft tumors from mice treated with or without losartan. **(A)** Normalized enrichment scores indicate the distribution of Gene Ontology categories. N=3 mice/ea. **(B)** GSEA enrichment plots. FDR <0.05. In the BR5 syngeneic OvCa model, flow cytometry analysis of **(C)** peritoneal tumors and **(D)** ascites collected on day 35 post-implantation. In the ID8 syngeneic OvCa model, flow cytometry analysis of **(E)** peritoneal tumors and **(F)** ascites collected on day 35 post-implantation. N=8 mice/ea. Data presented as Mean ± SD. See Table S2 for antibody panels used for flow cytometry.

Motivated by this fibrogenesis-downregulated and chemokine-enriched signature, we analyzed the cellular infiltrates in OvCa peritoneal tumors and ascites. In the BR5 syngeneic OvCa model, we found that losartan treatment increased tumor infiltration of immune effector CD4 and CD8 T cells, NK cells, and increased CD8 T cell/T_reg_ ratio. Both immune suppressive T_reg_ and myeloid-derived suppressor cells (MDSCs) were reduced in losartan-treated tumors (Fig 1C). In the ascites from losartan-treated mice, the percentage of CD8 T cells was significantly higher, and the percentage of CD45^+^ myeloid cells that were macrophages was significantly lower as compared with control mice. More importantly, losartan treatment induced expansion of interferon (IFN)-γ-producing CD8^+^ T cell population in the ascites, indicating activation of CD8 T cells (Fig 1D). The losartan-induced increase in tumor-infiltrating CD4 and CD8 T cells was further confirmed by immunofluorescent analysis in the BR5 model (sFig 1A-B). In a second syngeneic ID8 OvCa model, which forms small solid tumors throughout the peritoneal surface and large amounts of ascites, losartan treatment increased CD8 T cells and NK cells in the peritoneal tumors (Fig 1E) and ascites (Fig 1F), activated CD8 T cells in the ascites as demonstrated by the increased IFNγ^+^CD8^+^ T cells, TNFα^+^CD8^+^ T cells, and an increase in the CD8 T cell/T_reg_ ratio (Fig 1F). These studies suggest that in addition to normalizing the ECM in the tumor microenvironment, losartan treatment could reprogram the immune microenvironment from immunosuppressive to immunostimulatory.

### Combined losartan treatment enhances αPD1 efficacy in OvCa models

Motivated by losartan’s ability to increase immune cell infiltration and activation in the tumor, we explored combination treatments pairing losartan with a checkpoint inhibitor, anti-PD1 (αPD1), in two syngeneic OvCa models (Fig 2A). In both BR5 (Fig 2B) and ID8 (Fig 2C) models, we found that: i) αPD1 treatment significantly reduced both peritoneal tumor burden and ascites; ii) losartan treatment alone did not affect tumor growth, but reduced ascites; and iii) when losartan was combined with αPD1, it significantly enhanced the anti-tumor effect of αPD1. In the ID8 model, combination therapy also demonstrated superior efficacy in reducing ascites volume compared to αPD1 monotherapy. In peritoneal tumors collected from mice in the combination treatment group, we observed more apoptotic tumor cells (TUNEL^+^) and fewer proliferating tumor cells (PCNA^+^) compared to those in the control or monotherapy groups (sFig 2A).

**Figure 2.**
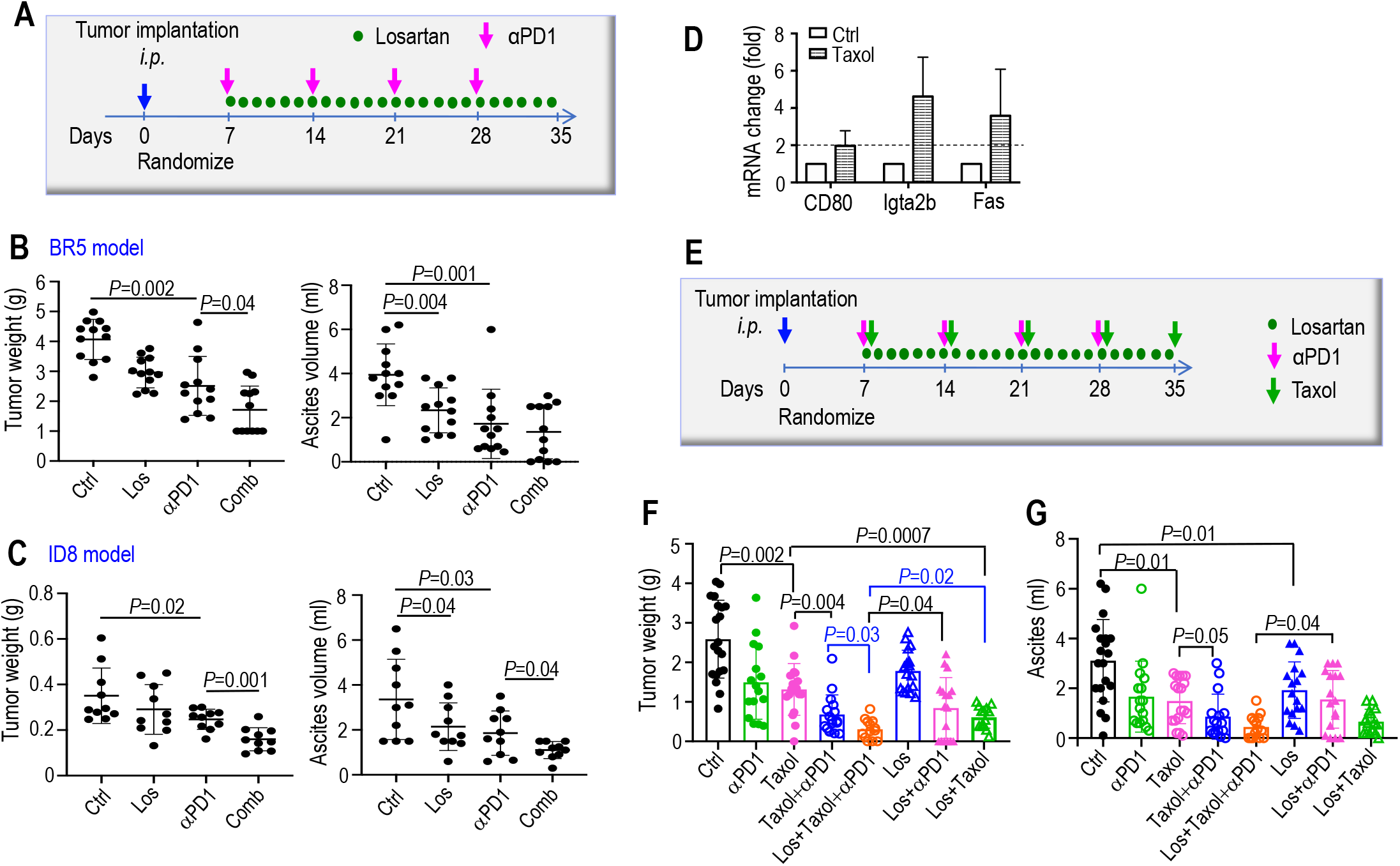
Combined losartan treatment improves chemo-immunotherapy efficacy in OvCa models. **(A)** Experimental design. Mice were injected *i*.*p*. with G-luc transduced OvCa tumor cells, and peritoneal tumor growth was monitored by peripheral blood G-luc levels. Between 7-10 days after implantation when blood G-luc value reached 2x10^6^ RLU, mice were randomized into treatment groups receiving: i) saline (control, ctrl), ii) losartan (Los, 40 mg/kg, QD), iii) αPD1 (200 μg/week for 4 weeks), v) losartan+αPD1. When mice became moribund, peritoneal tumor weight and ascites volume were measured in **(B)** BR5 model, Ctrl, Los, αPD1, and combination, n=12/ea; and **(C)** ID8 model, Ctrl (n=9), Los (n=10) αPD1 (n=10), and combination (n=10). Data presented are mean ± SEM. **(D)** BR5 tumors collected from mice treated with control (Ctrl) or Taxol (5 mg/kg, weekly) were lysed for RNA extraction. The expression of neoantigens was measured using qRT-PCR analysis. **(E)** Treatment schedule. Mice were injected *i*.*p*. with BR5 cells, and randomized as described in (A) into treatment groups receiving: i) saline (ctrl), n=20; ii) losartan, n=17; iii) Taxol, n=18; iv) αPD1, n=16; v) losartan+αPD1, n=18; vi) losartan+Taxol, n=15; vii) Taxol+αPD1, n=16; and viii) losartan+Taxol+αPD1, n=14. When mice became moribund, **(F)** peritoneal tumor weight, and **(G)** ascites volume were measured. Data presented are mean ± SEM. See the sequence of primers used in qRT-PCR in Table S3.

### Combined losartan treatment improves chemo-immunotherapy in OvCa models

Chemotherapy is the standard of care for OvCa patients. In the BR5 and ID8 OvCa models, Taxol treatment induced tumor cell death (sFig 2B) and generated neoantigens (Fig 2D), which play a critical role in mediating the response to immunotherapy ^27-29^. Previously, in two human ovarian cancer models, we demonstrated that losartan treatment increases intratumoral delivery of chemotherapy drugs ^25^. Based on these data, we evaluated whether losartan+αPD1+Taxol is superior to mono- and double combination therapies (Fig 2E). We found that combining losartan treatment: i) with Taxol, improves Taxol efficacy; this is consistent with our findings in the human OvCa models ^25^, ii) with αPD1, improves αPD1 efficacy, and iii) with Taxol+ αPD1, further improves the efficacy of chemo+immunotherapy by reducing tumor burden (Fig 2F) and ascites (Fig 2G), as compared to double combination treatments.

### Losartan treatment increases intratumoral αPD1 drug delivery

We investigated the mechanisms underlying the enhancement of chemo-immunotherapy by losartan. Previously we demonstrated that losartan treatment enhances chemotherapy efficacy via normalizing the ECM to improve blood vessel perfusion and chemotherapy drug delivery ^25^. To examine whether losartan affects the delivery of large molecular weight antibodies, we labeled the αPD1 antibody with a green fluorescein tag ^30^, injected it i.p. into control- or losartan-treated OvCa tumor-bearing mice, and collected peritoneal tumors 24 hours later. In losartan-treated tumors, we observed a significantly increased intratumoral fluorescence signal, indicating increased delivery of αPD1 antibody (Fig 3A-C).

**Figure 3.**
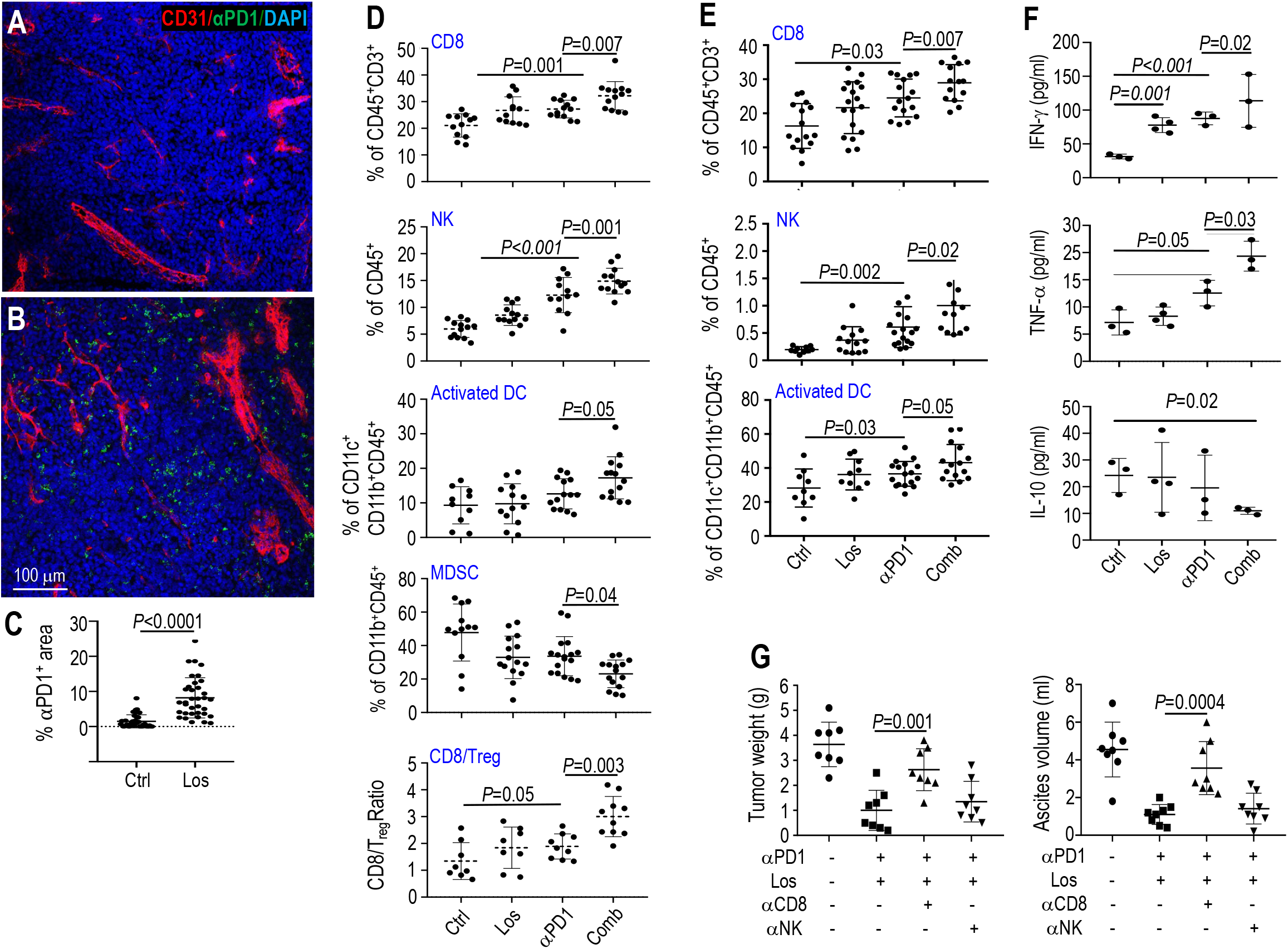
Losartan treatment enhances αPD1 efficacy, via increasing intratumoral drug delivery and immune effector cell infiltration in mouse OvCa models. Representative images of the intratumoral distribution of FITC-labeled αPD1 antibody (green) in **(A)** control and **(B)** losartan-treated tumors. Red, CD31^+^ tumor blood vessels; blue, DAPI. **(C)** Quantification of the fraction of tumor area positive for αPD1 antibody was performed using ImageJ. Flow cytometry analysis of immune cell composition in **(D)** peritoneal tumors and **(E)** ascites was performed 35 days after BR5 tumor implantation when significant differences in growth were evident. **(F)** Multiplex ELISA analysis of cytokine production in BR5 peritoneal tumors collected on day 35 after tumor implantation. Data presented are mean ± SD. **(G)** Depletion of CD8 cells (anti-CD8, 200 μg) or NK cells (anti-NK1.1, 25 μg) in BR5 model. Both antibodies were from BioXcell and administered *i*.*p*. once. Data presented are mean ± SEM.

### Losartan-enhanced αPD1 efficacy is mediated by CD8 T cells

Next, we characterized how the combination treatment changes intratumoral immune cell infiltration. Compared to the αPD1 monotherapy, combined losartan treatment resulted in a significant increase of tumor-infiltrating CD8 T cells, NK cells, and activated antigen-presenting dendritic cells (DCs), and a significant reduction of immune suppressive MDSCs. Moreover, the ratio of CD8/T_reg_ was further increased in the combination treatment group compared to αPD1 monotherapy (Fig 3D). Lymphocytic subpopulations in the ascites were also analyzed. Changes in CD8 T cells, NK cells, and activated DCs were consistent with those found in the tumor (Fig 3E), which was not the case for MDSCs and CD8/Treg ratio (sFig 3). Analyzing the peritoneal tumor secretome, we found that i) losartan treatment increased IFN-γ production, ii) αPD1 monotherapy increased both IFN-γ and TNF-α, and iii) combined losartan treatment not only further enhanced αPD1-induced IFN-γ and TNF-α production, but also significantly reduced immune suppressive IL-10 production (Fig 3F).

These data suggest that an influx of immune effector CD8 T cells and NK cells and activation of their immune cytokine production may be the major immunologic mechanism mediating the benefit of combined treatment. To test this hypothesis, we depleted mice of CD8 T cells or NK cells before treating them with αPD1 therapy. When the mice were depleted of CD8 T cells, the tumor control benefit of the combination therapy was abrogated, whereas depletion of NK cells had no effect (Fig 3G). These results suggest that losartan treatment enhances αPD1 efficacy via increasing CD8 T cells intratumoral infiltration.

### Losartan treatment suppresses IGF-1 signaling in OvCa model

To uncover potential losartan-activated molecular mechanisms that sensitize OvCa to chemo-immunotherapy, we profiled losartan-induced i) changes in the transcriptome by RNASeq, and ii) changes in signal transduction by receptor tyrosine kinase phosphorylation array. In the BR5 peritoneal tumors, losartan treatment suppressed the expression of genes in the Insulin Growth Factor (IGF)-1 pathway (Fig 4A-B), and reduced phosphorylation of both the IGF-1 receptor and Insulin receptor (Fig 4C-D).

**Figure 4.**
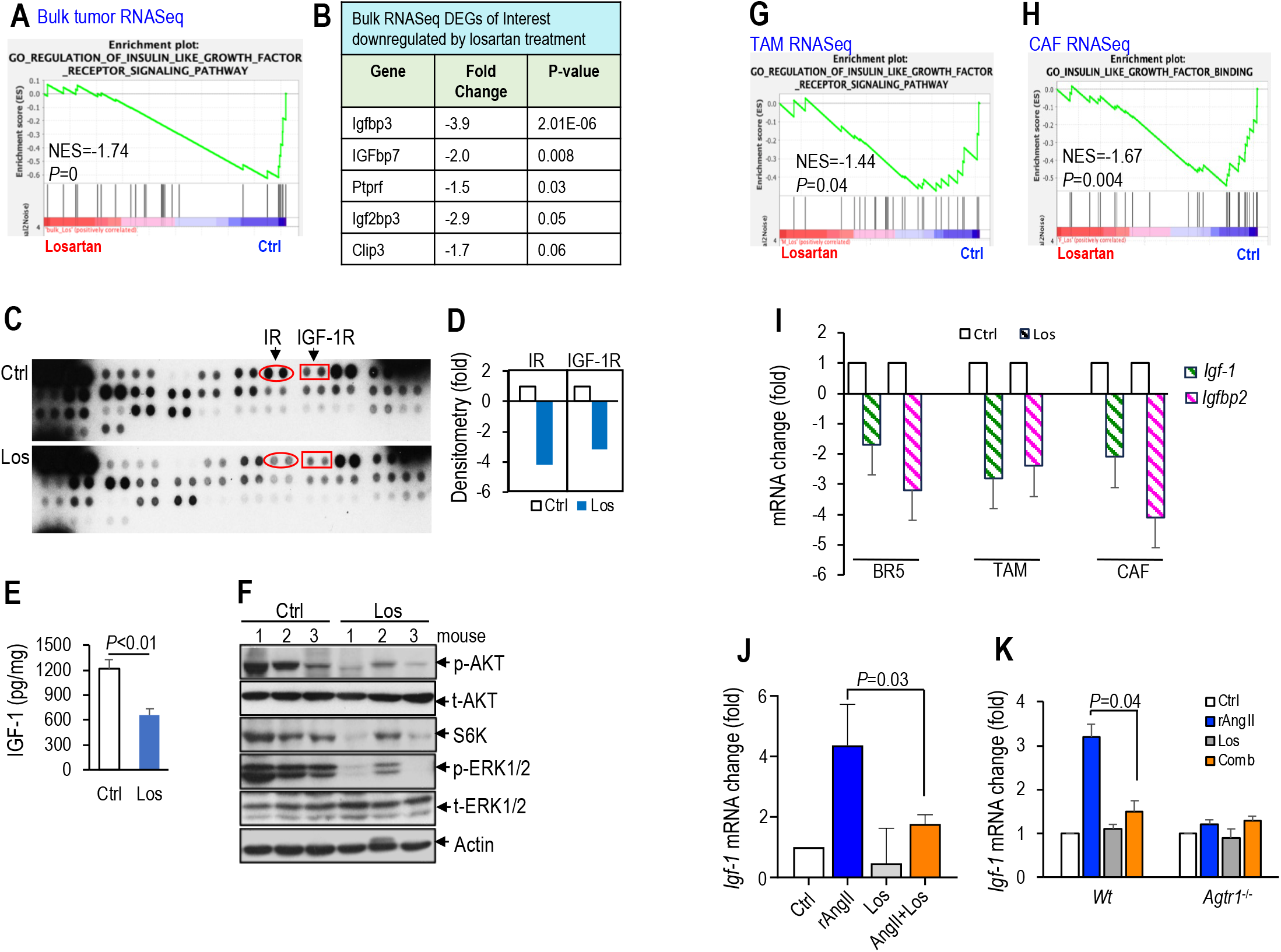
Losartan treatment suppresses IGF signaling in the OvCa model. Results of RNASeq analysis of BR5 OvCa peritoneal tumors from mice treated with control or losartan. **(A)** Gene Ontology enrichment plot demonstrating IGF receptor signaling downregulated in losartan-treated tumors. N=3 mice/ea. **(B)** Differentially expressed genes in the IGF pathway downregulated by losartan treatment. **(C)** Proteome array to screen for changes in receptor tyrosine kinase phosphorylation. Red oval: insulin receptor (IR), red rectangle: IGF-1R. **(D)** Densitometry analysis of IR and IGF-1R expression changes in losartan-treated tumors normalized to the control group. **(E)** murine IGF-1 protein level in ascites was measured by ELISA, N=3 mice/ea. **(F)** IGF-1 downstream AKT/S6 kinase and ERK1/2 MAPK phosphorylation were evaluated by western blot. GSEA analysis of RNASeq data of **(G)** TAMs and **(H)** CAFs isolated from peritoneal BR5 tumors. **(I)** Confirmation of *muIgf-1* and *muIgfbp2* mRNA levels change in BR5 tumor cells, TAMs, and CAFs isolated from control and losartan-treated BR5 tumors by qRT-PCR. N=3 tumors/group. In *in vitro* experiment, **(J)** BR5 cells and **(K)** peritoneal macrophages isolated from wildtype or *Agtr1*^-/-^ mice were treated with recombinant AngII (0.1 μM), losartan (1 μM) or AngII+Los for 6 hours. Changes in mouse *Igf-1* RNA were evaluated by qRT-PCR. Data presented are mean ± SD.

The IGF axis has been shown to play a pivotal role in the development, progression, and chemotherapy resistance of ovarian cancer ^31^. IGF-1 is the key member of the IGF family that drives tumor progression. We confirmed the above transcriptional results by demonstrating that losartan treatment reduced the secretion of IGF-1 protein into the ascites (Fig 4E) and reduced IGF-1 downstream AKT/S6 kinase and ERK1/2 MAPK signaling in the peritoneal tumors (Fig 4F). OvCa tumor cells express dysregulated IGF-1 pathway genes ^31^, and tumor-associated macrophages (TAMs) and cancer-associated fibroblasts (CAFs) have been shown to be significant contributors to IGF-1 production in invasive breast cancer ^32^. To pinpoint the cell type that responds to the IGF-regulating effects of losartan, we isolated TAMs and CAFs from the BR5 peritoneal tumors and performed RNASeq. GSEA analysis of the RNASeq data revealed that losartan exhibited a suppressive effect on IGF-1R signaling in both TAMs and CAFs (Fig 4G-H, Table S4,S5). We confirmed the profiling data by showing that losartan treatment decreased mRNA levels of IGF-1, as well as IGF Binding Protein 2 (IGF-BP2), a regulator of IGF-1 availability and function through binding, in BR5 tumor cells, as well as in TAMs and CAFs (Fig 4I).

To investigate if losartan plays a causal role in the regulation of IGF-1 expression, we treated BR5 tumor cells in culture with recombinant angiopoietin II (Ang II), losartan, or a combination of AngII+losartan. We observed that losartan treatment blocked AngII-induced IGF-1 expression (Fig 4J). Next, we investigate if losartan regulates IGF-1 expression by blocking the AngII receptor (AT1). To this end, we isolated peritoneal macrophages from wildtype (*Wt*) and from AT1 knockout mice (*Agtr1*^*-/-*^). In macrophages from *Wt* mice, AngII induced IGF-1 mRNA expression, and the induction was abolished by losartan treatment. Conversely, in *Agtr1*^*-/-*^ mice macrophages, AngII failed to induce IGF-1 mRNA expression (Fig 4K). These findings suggest that losartan can directly downregulate IGF-1 expression in OvCa cancer cells and TAMs by blocking the AT1 receptor.

### Losartan treatment, via suppressing the IGF-1 signaling, augments chemo-immunotherapy

To investigate if the IGF-1 reduction is essential in mediating the losartan-enhanced treatment sensitivity, we stably overexpressed IGF-1 in the BR5 cells. *In vitro* experiments showed that IGF-1 overexpression had no effect on cancer cell proliferation (sFig 4A). Similarly, in the mouse model, IGF-1 overexpression did not change peritoneal tumor growth or ascites formation (sFig 4B-C). Our mechanistic study further confirmed that IGF-1 overexpression does not affect tumor cell proliferation or apoptosis (sFig 4D-E). IGF signaling activation contributes to resistance to chemotherapy, endocrine therapy, radiotherapy, and targeted therapy in different types of cancers ^33^. Our study revealed that overexpression of IGF-1 in BR5 tumor cells led to resistance to Taxol treatment *in vitro* (Fig 5A), and caused BR5 peritoneal tumors to become refractory to Taxol treatment in mouse models (Fig 5B-D).

**Figure 5.**
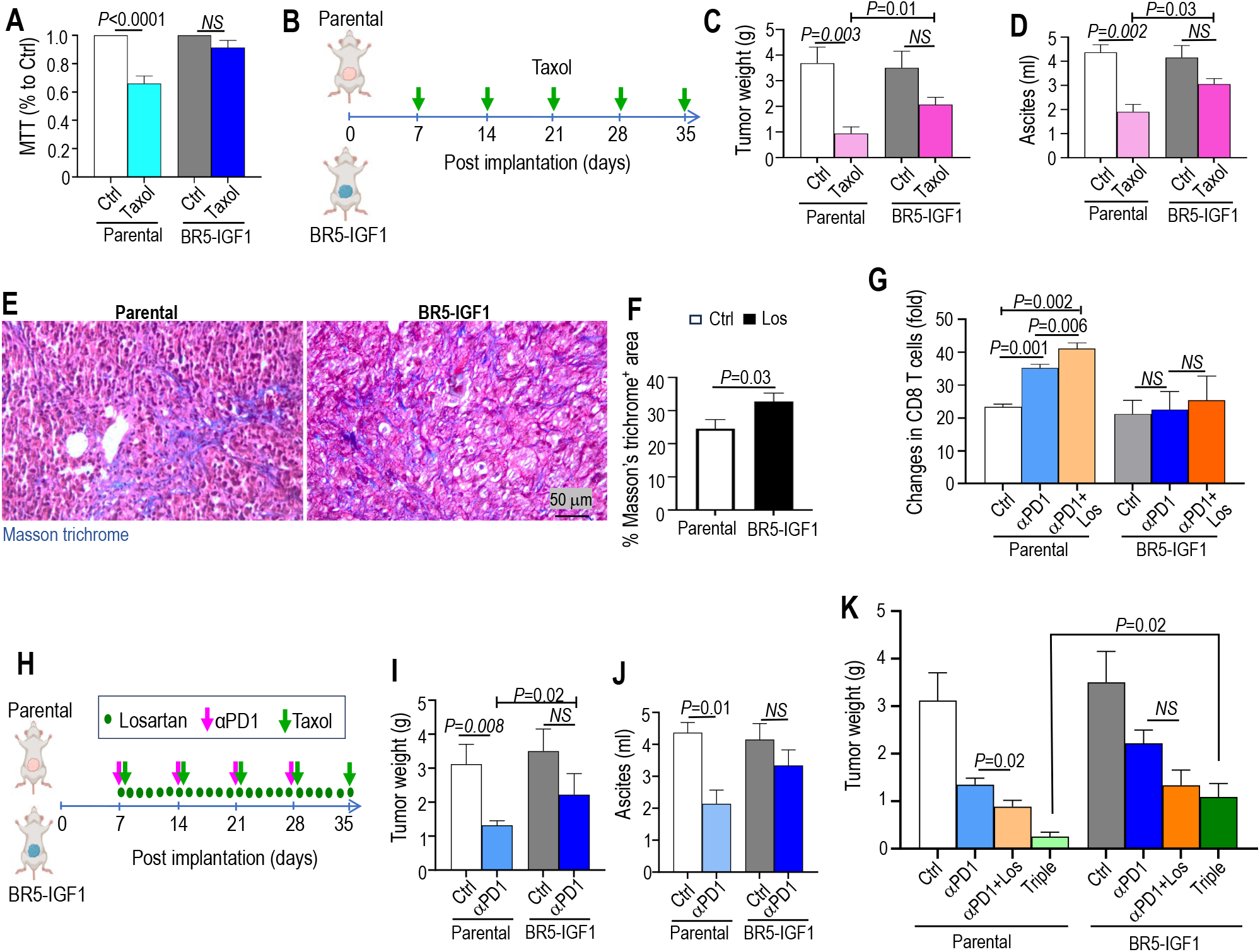
Losartan treatment, via suppressing the IGF-1 signaling, enhances chemo-immunotherapy. **(A)** Parental and IGF-1 overexpressing BR5 cells were plated in 96-well plate (1x10^4^ cells/well) and treated with DMEM (Ctrl) or Taxol (10 nM) for 48 hours. Cell viability was evaluated by MTT assay. **(B)** Experimental design. Mice were injected *i*.*p*. with parental, mock- or IGF-1 transfected BR5 OvCa tumor cells, and randomized into treatment groups receiving control (saline) or Taxol (4 mg/kg, weekly), n=6/ea. When mice became moribund, **(C)** peritoneal tumor weight and **(D)** volume of ascites were measured. **(E)** Masson’s Trichrome staining of parental and IGF-1 transfected BR5 peritoneal tumors. **(F)** The percentage of Masson’s Trichrome positive staining area was quantified using Image J. **(G)** peritoneal tumors were collected for flow cytometry to profile changes in CD8 T cells. **(H-J)** Mice bearing parental or IGF-1 transfected tumors were treated with control IgG (Ctrl) or αPD1, when mice became moribund **(I)** tumor weight, and **(J)** ascites volume was measured. **(K)** A different cohort of mice bearing parental or IGF-1 transfected tumors was treated with control IgG (Ctrl), αPD1, αPD1+losartan, or triple combination therapy of αPD1+losartan+Taxol (Triple). N=6/each, data presented are mean ± SEM.

In IGF-1 overexpressing BR5 tumors, we observed increased collagen levels in the tumor matrix (Fig 5E-F). In tumors, the abnormal collagen density and alignment can limit T cell motility, and tumor cell killing ^34^. In IGF-1 overexpressing BR5 tumors, we found αPD1 treatment failed to induce intratumoral infiltration of CD8 T cells, and combined losartan treatment failed to further enhance the αPD1-induced CD8 T cell recruitment (Fig 5G). Consequently, we observed that overexpression of IGF-1 abolished the inhibitory effects of αPD1 on tumor growth and ascites formation (Fig 5H-J). As IGF-1 overexpression causes both chemo- and immunotherapy resistance, IGF overexpression abolished the benefit of losartan in enhancing the efficacy of combined Taxol and αPD1 treatment (Fig 5K). Collectively, these data suggest that the IGF-1 axis contributes to OvCa resistance to αPD1 and chemotherapy, and is critical in mediating the effects of losartan in enhancing chemo-immunotherapy efficacy.

### Losartan treatment is associated with reduced matrix level, fibrogenic signaling, and increased immune cell infiltration in OvCa patients

To validate our preclinical findings in OvCa patients, we obtained paired snap-frozen RNA samples and paraffin-embedded archived OvCa tumor tissues collected from patients who had been treated with or without losartan (Table S6). RNASeq analysis of the patient RNA samples demonstrated that losartan treatment down-regulated genes in the renin-angiotensin system pathways, ECM, and fibrogenic signaling, such as NOTCH, TGF-β, Hedgehog, MAPK, and Focal adhesion pathways (Fig 6A). GSEA analysis of the differentially regulated genes demonstrated that i) downregulated genes are enriched in the ECM and fibrogenic TGF-β signaling pathways; and ii) upregulated genes are enriched in the innate immune response and oxidative phosphorylation, which plays a significant role in regulating the immune response (Fig 6B). In paraffin-embedded patient tumors, we found that treatment with losartan is associated with lower collagen levels (Fig 6C), more tumor-infiltrating CD4 and CD8 T cells (Fig 6D-E), and lower IGF-1 levels (Fig 6F), as compared to those without losartan treatment.

**Figure 6.**
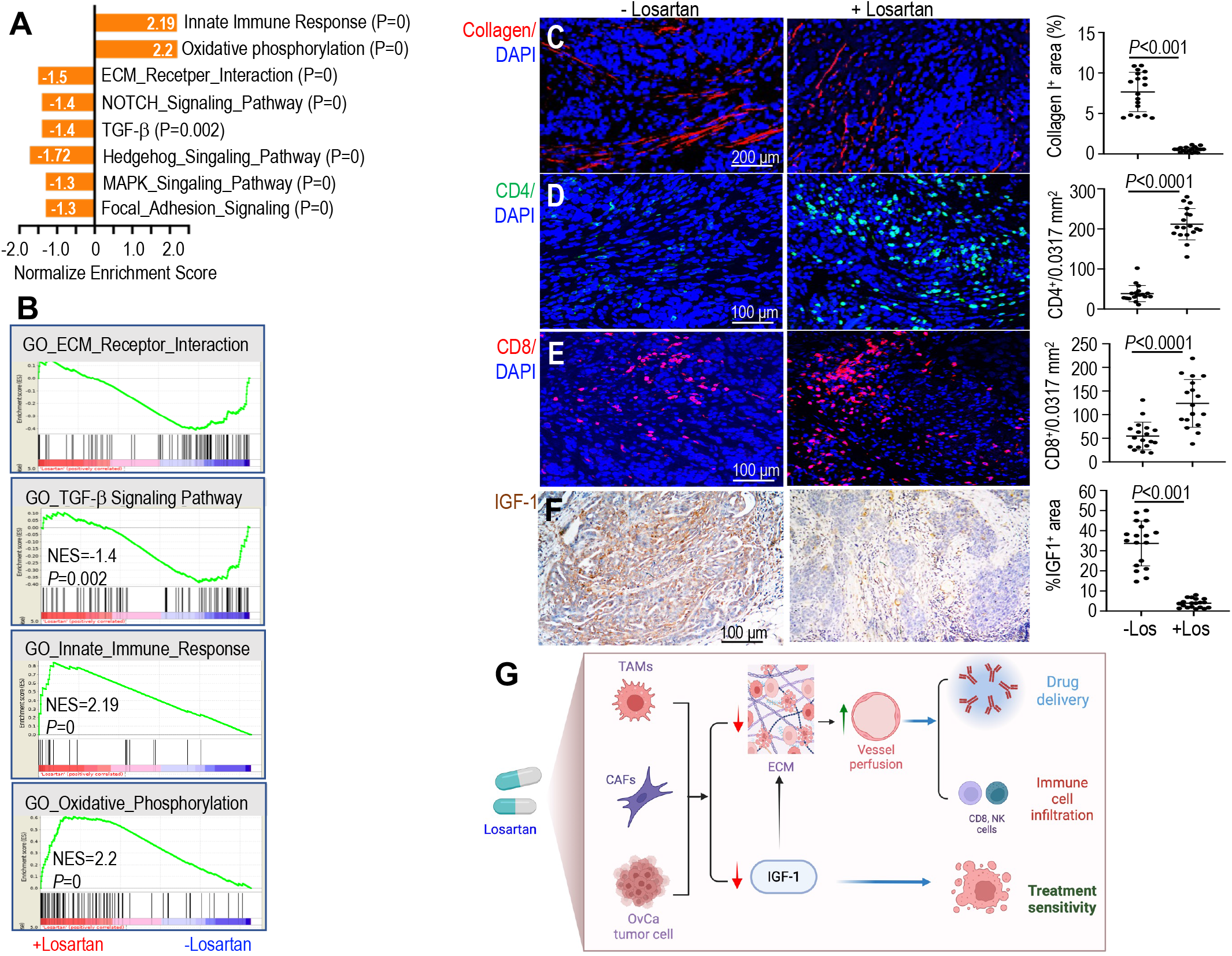
Losartan treatment is associated with reduced matrix level, fibrogenic signaling, and increased immune cell infiltration in OvCa patients. Snap-frozen OvCa patient tumor samples were subject to RNASeq analysis. **(A)** Normalized enrichment scores indicate the distribution of Gene Ontology categories. N=3 patients/each arm. **(B)** GSEA enrichment plots. FDR <0.05. Archived paraffin-embedded tumors from patients with OvCa and treated without (n=3) or with losartan (n=3) were stained for **(C)** collagen I (red). The fractions of Collagen I (red) positive areas were digitally quantified using ImageJ software. **(D)** CD4 T cells (green), and **(E)** CD8 T cells (red). The number of CD4 and CD8 T cells was manually counted. DAPI (blue). **(F)** representative IGF-1 IHC staining images. The fractions of IGF-1 (brown) positive areas were digitally quantified using ImageJ software. Images of 20 random fields were analyzed, and data presented as mean ± SD. **(G)** Schematics of losartan mechanism of action. In OvCa models, losartan exhibits dual effects on both the tumor microenvironment and cancer cells: i) on the TME, losartan treatment reduces matrix content leading to increased vascular perfusion, and thus enhances ICI antibody delivery and intratumoral infiltration and function of immune effector cells; and ii) on the OvCa cells, losartan treatment suppresses the IGF-1 signaling, resulting in enhanced chemosensitivity. With its dual impact on both tumor cells and the TME, losartan treatment enhances the efficacy of chemo-immunotherapy in OvCa models.

## Discussion

OvCa has the highest mortality rate of all gynecological cancers. Despite high initial response rates towards chemotherapy, approximately 70% of all patients relapse and develop resistant disease within five years after diagnosis ^2^. There is an urgent need for the development of novel and effective therapeutic strategies. Immunotherapies that are efficacious in the treatment of many malignant neoplasms have shown limited efficacy in OvCa to date. The JAVELIN Ovarian 200 RCT trial of PD-L1 inhibitor, avelumab ^35^, and the NINJA trial of PD-1 inhibitor, nivolumab ^36^, did not demonstrate improved progression-free survival (PFS) or overall survival (OS). These studies highlight the need to develop strategies to increase immune infiltrates in tumors.

Here, using OvCa models, we discovered that losartan exhibits dual effects on both the tumor microenvironment and cancer cells: i) on the TME, losartan treatment reduces matrix content leading to increased vascular perfusion, and thus enhances ICI antibody delivery and immune effector cell intratumoral infiltration and function; and ii) on the OvCa cells, losartan treatment suppresses the IGF-1 signaling, resulting in enhanced chemosensitivity. With its dual impact on both tumor cells and the TME, losartan treatment enhances the efficacy of chemo-immunotherapy in OvCa models (Fig 6G).

The density and spatial distribution of immune infiltrates in tumors are associated with patient survival and response to immune therapy in melanoma, pancreatic, and other cancers ^2,37-39^. In OvCa, although checkpoint inhibitors have had limited efficacy, a positive trend in objective response rate (ORR) was reported for patients with high CD8 T cells who received avelumab in JAVELIN Ovarian 200 ^35^ or pembrolizumab in KEYNOTE-100 ^40^. Multiple mechanisms impede the intratumoral infiltration of immune cells, such as abnormal vascular perfusion and ECM barriers. Decades of research have demonstrated that the OvCa TME is significantly abnormal, displaying a desmoplastic, highly fibrotic extracellular matrix with high levels of collagen. ECM is a major determinant of tissue architecture, which determines the direction, and efficiency of T cell movement ^41^. The high-density ECM generates solid stress that compresses blood vessels and impedes drug delivery in tumors ^11^. Previously, we discovered that losartan, an FDA-approved, widely prescribed angiotensin blocker to control hypertension, is capable of i)”normalizing” the extracellular matrix, ii) opening compressed tumor vessels, and iii) improving the delivery and efficacy of chemotherapy in OvCa ^25^. As a natural extension of this study, we investigated if losartan could improve intratumoral immune cell infiltration and thus reprogram the immunosuppressive TME of OvCa to enhance immunotherapy. Here, using two syngeneic immunocompetent mouse OvCa models, we demonstrated that losartan treatment i) improved ICI antibody delivery, ii) increased immune cell intratumoral infiltration and function, and as a result, iii) enhanced the efficacy of chemo-immunotherapy in ovarian cancer models.

In addition to normalizing the TME, we found that losartan treatment, via directly regulating IGF-1 signaling in both cancer and stromal cells, enhances OvCa sensitivity to chemo-immunotherapy. The IGF system is composed of ligands (IGF-1, IGF-2), IGF receptors (IGF-1R, IGF-2R), and six IGF binding proteins (IGFBPs 1-6) ^34^. Elevated activation of the IGF signaling pathway is implicated in the development, progression, and treatment resistance of many types of cancer ^31,42-45^. Activation of the IGF receptors mediates key processes in cancer, such as cell growth, angiogenesis, and DNA damage repair ^46^. In ovarian cancers, dysregulated IGF-1 family gene expression has been reported to correlate with disease progression and resistance to platinum-based chemotherapy ^31,42^. Here in OvCa models, we found that losartan treatment reduced IGF-1 secretion, IGF-1R activation, and IGF downstream AKT and ERK1/2 MAPK signaling. In our study, we further explored the cellular and molecular mechanisms by which losartan regulates IGF-1 signaling. Crosstalk between IGF-1 and AngII pathways has been described in the cardiovascular system - AngII stimulates IGF-1 transcription in vascular smooth muscle cells ^47^, and AngII blockade with angiotensin-converting enzyme (ACE) inhibitor captopril and with AT1 blocker losartan reduced IGF-1 binding to its receptors in a rat model of cardiac hypertrophy ^48^. However, the crosstalk between IGF-AngII pathways has not been reported in cancer. We discovered that losartan downregulated IGF-1 pathway genes in both OvCa tumor cells and tumor stromal cells (TAMs, and CAFs). In cancer cells and macrophages, AngII directly induced IGF-1 expression, and blocking the AT1 receptor with losartan abolished this AngII induction of IGF-1. Using *Agtr1*^*-/-*^ mice, we further demonstrated that AngII directly regulates IGF-1 mRNA expression via the AT1 receptor. In addition to regulating the IGF-1 ligand expression, losartan also reduces the expression of IGFBP2, a molecule initially discovered to modulate the bioavailability of IGF-1 through binding, and is now known to activate pro-survival and pro-invasive ERK and Src signaling in tumors ^49,50^. In the clinic, high levels of IGFBP2 have been associated with poor survival in OvCa ^51^. In our study, using tumors from patients with OvCa that have been treated with or without losartan, we confirmed that losartan treatment is associated with a reduced IGF-1 level in human tumors. Our study revealed the heretofore unknown cellular and molecular mechanisms of how Angiotensin signaling regulates the IGF-1 pathway in OvCa tumors.

Next, we investigated the functional role of the IGF-1 axis in OvCa tumor progression and treatment response, as well as its role in mediating the losartan effects. We demonstrated that overexpression of IGF-1 did not affect OvCa tumor progression, but rendered tumors insensitive to chemotherapy. This is consistent with published findings that IGF-1 signaling contributes to resistance to chemotherapy and targeted therapies ^52-54^. A recent study in syngeneic lung cancer models showed that short-term starvation sensitizes tumors to αPD1 treatment, and the enhanced anti-tumor activity was linked to a reduction of the IGF-1 level ^55^. Motivated by this report, we investigated if IGF-1 affects tumor immunity and response to immunotherapy in OvCa models. Based on our observation of an elevated matrix level in IGF-1 overexpressing tumors, coupled with the knowledge that ECM plays a pivotal role in T cell movement, we compared the immune effector cell infiltration in parental vs. IGF-1 overexpressing tumors. Here, we demonstrated that IGF-1 overexpression abolished αPD1-induced CD8 T cell and NK cell infiltration, consequently attenuating the efficacy of αPD1 treatment. Taken together, our data indicate that targeting the IGF-1 signaling holds therapeutic potential for OvCa, and supports the clinical investigation of IGF-1 inhibitors as well as losartan in combination with chemo-immunotherapy in OvCa patients.

## Materials and Methods

Tumor growth and treatment efficacy were investigated in two syngeneic mouse models of OvCa. For additional information regarding cell lines, animal models, treatment protocols, patient characteristics, and statistical analysis, see Supplemental Information.

## Supplementary Materials

### Materials and Methods

Figure S1. Histological confirmation of losartan-induced tumor-infiltrating CD4 and CD8 T cells.

Figure S2. Combined losartan treatment improves immunotherapy efficacy in OvCa models.

Figure S3. Losartan treatment’s effects on immune cell infiltration in mouse OvCa models.

Figure S4. IGF-1 overexpression does not affect OvCa tumor growth.

Table S1. Differentially expressed genes in human OvCa tumors and in mouse stromal cells. Table S2. Antibody panels used for flow cytometry.

Table S3. Primer sequences used in qRT-PCR assay.

Table S4. Differentially expressed genes in TAMs isolated from murine OvCa tumors treated with or without losartan.

Table S5. Differentially expressed genes in CAFs isolated from murine OvCa tumors treated with or without losartan.

Table S6. Demographic and clinical characteristics of OvCa patient samples.

## Notes

## Acknowledgments

We thank Dr. Brian Seed for providing the cloning vector, we thank Mark Duquette, Naifang Lu, and Anna Khachatryan for their superb technical support, and Dr. Peigen Huang for assisting in animal studies.

## Author contributions

L.X. and R.K.J. designed the research; Y.S., Z.Y., Y.Z., and S.L. performed mouse model studies; Y.S., Y.Z., C.Y., and S.L. performed histological studies; S.S. and P.L. analyzed RNASeq data; Y.S., L.W., B.R.R., and A.M. performed patient sample analysis; L.X., Y.S., Z.Y., L.W., S.S., P.L., A.M., L.Z. analyzed data; L.X. and R.K.J. wrote the paper.

## Funding

This study was supported by the American Cancer Society Mission Boost Award (to L.X.), NIH R01-NS126187 and R01-DC020724 (to L.X.), Department of Defense Investigator-Initiated Research Award (W81XWH-20-1-0222, to L.X.) and Clinical Trial Award (W81XWH2210439, to L.X.), Children’s Tumor Foundation Drug Discovery Initiative (to L.X.); NIH grant R35-CA197743, and in part through grants R01-R01CA259253, R01-CA269672, R01-NS118929, U01-CA224348 and U01-CA261842 and by Nile Albright Research Foundation, the National Foundation for Cancer Research, Harvard Ludwig Cancer Center, and Jane’s Trust Foundation (to R.K.J.); and Nile Albright Research Foundation, Vincent Memorial Hospital Foundation, NCI P50CA240243, Julie Fund, Worden Family Foundation (to B.R.R).

## Conflict of interest

R.K.J. received consultant fees from Cur, Elpis, Innocoll, SPARC, and SynDevRx; owns equity in Accurius, Enlight, and SynDevRx; is on the Board of Trustees of Tekla Healthcare Investors, Tekla Life Sciences Investors, Tekla Healthcare Opportunities Fund, and Tekla World Healthcare Fund; and received research grants from Boehringer Ingelheim and Sanofi. No funding or reagents from these companies were used in this study. B.R.R. reports serving on the advisory board for VincenTech which has no direct connection to the current research. The other authors have no competing interests to declare.

## Notes

### Competing Interest Statement

The authors have declared no competing interest.

## Reference

1 Siegel, R. L., Miller, K. D., Wagle, N. S. & Jemal, A. Cancer statistics, 2023. CA Cancer J Clin 73, 17–48, doi:10.3322/caac.21763 (2023).

2 Richardson, D. L., Eskander, R. N. & O’Malley, D. M. Advances in Ovarian Cancer Care and Unmet Treatment Needs for Patients With Platinum Resistance: A Narrative Review. JAMA Oncol, doi:10.1001/jamaoncol.2023.0197 (2023).

3 Gaillard, S. L., Secord, A. A. & Monk, B. The role of immune checkpoint inhibition in the treatment of ovarian cancer. Gynecol Oncol Res Pract 3, 11, doi:10.1186/s40661-016-0033-6 (2016).

4 Castellano, T., Moore, K. N. & Holman, L. L. An Overview of Immune Checkpoint Inhibitors in Gynecologic Cancers. Clin Ther 40, 372–388, doi:10.1016/j.clinthera.2018.01.005 (2018).

5 Bak, S. P., Alonso, A., Turk, M. J. & Berwin, B. Murine ovarian cancer vascular leukocytes require arginase-1 activity for T cell suppression. Mol Immunol 46, 258–268, doi:10.1016/j.molimm.2008.08.266 (2008).

6 Curiel, T. J. et al. Specific recruitment of regulatory T cells in ovarian carcinoma fosters immune privilege and predicts reduced survival. Nat Med 10, 942–949, doi:10.1038/nm1093 (2004).

7 Wolf, D. et al. The expression of the regulatory T cell-specific forkhead box transcription factor FoxP3 is associated with poor prognosis in ovarian cancer. Clin Cancer Res 11, 8326–8331, doi:10.1158/1078-0432.CCR-05-1244 (2005).

8 Samrao, D. et al. Histologic parameters predictive of disease outcome in women with advanced stage ovarian carcinoma treated with neoadjuvant chemotherapy. Transl Oncol 5, 469–474 (2012).

9 Maniati, E. et al. Mouse Ovarian Cancer Models Recapitulate the Human Tumor Microenvironment and Patient Response to Treatment. Cell Rep 30, 525–540 e527, doi:10.1016/j.celrep.2019.12.034 (2020).

10 Chauhan, V. P. et al. Normalization of tumour blood vessels improves the delivery of nanomedicines in a size-dependent manner. Nat Nanotechnol 7, 383–388, doi:10.1038/nnano.2012.45 (2012).

11 Jain, R. K., Martin, J. D. & Stylianopoulos, T. The role of mechanical forces in tumor growth and therapy. Annu Rev Biomed Eng 16, 321–346, doi:10.1146/annurev-bioeng-071813-105259 (2014).

12 Stylianopoulos, T., Munn, L. L. & Jain, R. K. Reengineering the Physical Microenvironment of Tumors to Improve Drug Delivery and Efficacy: From Mathematical Modeling to Bench to Bedside. Trends Cancer 4, 292–319, doi:10.1016/j.trecan.2018.02.005 (2018).

13 Clever, D. et al. Oxygen Sensing by T Cells Establishes an Immunologically Tolerant Metastatic Niche. Cell 166, 1117–1131 e1114, doi:10.1016/j.cell.2016.07.032 (2016).

14 Fukumura, D., Kloepper, J., Amoozgar, Z., Duda, D. G. & Jain, R. K. Enhancing cancer immunotherapy using antiangiogenics: opportunities and challenges. Nature Reviews Clinical Oncology 15, 325–340 (2018).

15 Huang, Y. et al. Vascular normalizing doses of antiangiogenic treatment reprogram the immunosuppressive tumor microenvironment and enhance immunotherapy. Proc Natl Acad Sci U S A 109, 17561–17566, doi:10.1073/pnas.1215397109 (2012).

16 Jain, R. K. Antiangiogenesis strategies revisited: from starving tumors to alleviating hypoxia. Cancer Cell 26, 605–622, doi:10.1016/j.ccell.2014.10.006 (2014).

17 Manning, E. A. et al. A vascular endothelial growth factor receptor-2 inhibitor enhances antitumor immunity through an immune-based mechanism. Clin Cancer Res 13, 3951–3959, doi:10.1158/1078-0432.CCR-07-0374 (2007).

18 Martin, J. D., Fukumura, D., Duda, D. G., Boucher, Y. & Jain, R. K. Reengineering the Tumor Microenvironment to Alleviate Hypoxia and Overcome Cancer Heterogeneity. Cold Spring Harb Perspect Med 6, a027094, doi:10.1101/cshperspect.a027094 (2016).

19 Ruffell, B. & Coussens, L. M. Macrophages and therapeutic resistance in cancer. Cancer Cell 27, 462–472, doi:10.1016/j.ccell.2015.02.015 (2015).

20 Munn, L. L. & Jain, R. K. Vascular regulation of antitumor immunity. Science 365, 544–545, doi:10.1126/science.aaw7875 (2019).

21 Chauhan, V. P. et al. Angiotensin inhibition enhances drug delivery and potentiates chemotherapy by decompressing tumour blood vessels. Nat Commun 4, 2516–2527, doi:10.1038/ncomms3516 (2013).

22 Diop-Frimpong, B., Chauhan, V. P., Krane, S., Boucher, Y. & Jain, R. K. Losartan inhibits collagen I synthesis and improves the distribution and efficacy of nanotherapeutics in tumors. Proc Natl Acad Sci U S A 108, 2909–2914, doi:10.1073/pnas.1018892108 (2011).

23 Pinter, M. & Jain, R. K. Targeting the renin-angiotensin system to improve cancer treatment: Implications for immunotherapy. Sci Transl Med 9, eaan5616, doi:10.1126/scitranslmed.aan5616 (2017).

24 Regan, D. P. et al. Losartan Blocks Osteosarcoma-Elicited Monocyte Recruitment, and Combined With the Kinase Inhibitor Toceranib, Exerts Significant Clinical Benefit in Canine Metastatic Osteosarcoma. Clin Cancer Res 28, 662–676, doi:10.1158/1078-0432.CCR-21-2105 (2022).

25 Zhao, Y. et al. Losartan treatment enhances chemotherapy efficacy and reduces ascites in ovarian cancer models by normalizing the tumor stroma. Proc Natl Acad Sci U S A, pii:201818357. 201818310.201811073/pnas.1818357116, doi:10.1073/pnas.1818357116 (2019).

26 Cen, X., Liu, S. & Cheng, K. The Role of Toll-Like Receptor in Inflammation and Tumor Immunity. Front Pharmacol 9, 878, doi:10.3389/fphar.2018.00878 (2018).

27 van Rooij, N. et al. Tumor exome analysis reveals neoantigen-specific T-cell reactivity in an ipilimumab-responsive melanoma. J Clin Oncol 31, e439–442, doi:10.1200/JCO.2012.47.7521 (2013).

28 Robbins, P. F. et al. Mining exomic sequencing data to identify mutated antigens recognized by adoptively transferred tumor-reactive T cells. Nat Med 19, 747–752, doi:10.1038/nm.3161 (2013).

29 Gubin, M. M. et al. Checkpoint blockade cancer immunotherapy targets tumour-specific mutant antigens. Nature 515, 577–581, doi:10.1038/nature13988 (2014).

30 Zhdanov, D. D. et al. Murine regulatory T cells induce death of effector T, B, and NK lymphocytes through a contact-independent mechanism involving telomerase suppression and telomere-associated senescence. Cell Immunol 331, 146–160, doi:10.1016/j.cellimm.2018.06.008 (2018).

31 Liefers-Visser, J. A. L., Meijering, R. A. M., Reyners, A. K. L., van der Zee, A. G. J. & de Jong, S. IGF system targeted therapy: Therapeutic opportunities for ovarian cancer. Cancer Treat Rev 60, 90–99, doi:10.1016/j.ctrv.2017.08.012 (2017).

32 Ireland, L. et al. Blockade of insulin-like growth factors increases efficacy of paclitaxel in metastatic breast cancer. Oncogene 37, 2022–2036, doi:10.1038/s41388-017-0115-x (2018).

33 Grimberg, A. Mechanisms by which IGF-I may promote cancer. Cancer Biol Ther 2, 630–635 (2003).

34 Yahya, M. A., Sharon, S. M., Hantisteanu, S., Hallak, M. & Bruchim, I. The Role of the Insulin-Like Growth Factor 1 Pathway in Immune Tumor Microenvironment and Its Clinical Ramifications in Gynecologic Malignancies. Front Endocrinol (Lausanne) 9, 297, doi:10.3389/fendo.2018.00297 (2018).

35 Pujade-Lauraine, E. et al. Avelumab alone or in combination with chemotherapy versus chemotherapy alone in platinum-resistant or platinum-refractory ovarian cancer (JAVELIN Ovarian 200): an open-label, three-arm, randomised, phase 3 study. Lancet Oncol 22, 1034–1046, doi:10.1016/S1470-2045(21)00216-3 (2021).

36 Hamanishi, J. et al. Nivolumab Versus Gemcitabine or Pegylated Liposomal Doxorubicin for Patients With Platinum-Resistant Ovarian Cancer: Open-Label, Randomized Trial in Japan (NINJA). J Clin Oncol 39, 3671–3681, doi:10.1200/JCO.21.00334 (2021).

37 Erdag, G. et al. Immunotype and immunohistologic characteristics of tumor-infiltrating immune cells are associated with clinical outcome in metastatic melanoma. Cancer Res 72, 1070–1080, doi:10.1158/0008-5472.CAN-11-3218 (2012).

38 Carstens, J. L. et al. Spatial computation of intratumoral T cells correlates with survival of patients with pancreatic cancer. Nat Commun 8, 15095, doi:10.1038/ncomms15095 (2017).

39 Palucka, A. K. & Coussens, L. M. The Basis of Oncoimmunology. Cell 164, 1233–1247, doi:10.1016/j.cell.2016.01.049 (2016).

40 Matulonis, U. A. et al. Antitumor activity and safety of pembrolizumab in patients with advanced recurrent ovarian cancer: results from the phase II KEYNOTE-100 study. Ann Oncol 30, 1080–1087, doi:10.1093/annonc/mdz135 (2019).

41 Hallmann, R. et al. The regulation of immune cell trafficking by the extracellular matrix. Curr Opin Cell Biol 36, 54–61, doi:10.1016/j.ceb.2015.06.006 (2015).

42 Yee, D., Morales, F. R., Hamilton, T. C. & Von Hoff, D. D. Expression of insulin-like growth factor I, its binding proteins, and its receptor in ovarian cancer. Cancer Res 51, 5107–5112 (1991).

43 Resnicoff, M., Ambrose, D., Coppola, D. & Rubin, R. Insulin-like growth factor-1 and its receptor mediate the autocrine proliferation of human ovarian carcinoma cell lines. Lab Invest 69, 756–760 (1993).

44 Ouban, A., Muraca, P., Yeatman, T. & Coppola, D. Expression and distribution of insulin-like growth factor-1 receptor in human carcinomas. Hum Pathol 34, 803–808, doi:10.1016/s0046-8177(03)00291-0 (2003).

45 Bruchim, I. & Werner, H. Targeting IGF-1 signaling pathways in gynecologic malignancies. Expert Opin Ther Targets 17, 307–320, doi:10.1517/14728222.2013.749863 (2013).

46 Werner, H. & LeRoith, D. The role of the insulin-like growth factor system in human cancer. Adv Cancer Res 68, 183–223, doi:10.1016/s0065-230x(08)60354-1 (1996).

47 Ma, Y. et al. Angiotensin II stimulates transcription of insulin-like growth factor I receptor in vascular smooth muscle cells: role of nuclear factor-kappaB. Endocrinology 147, 1256–1263, doi:10.1210/en.2005-0888 (2006).

48 Haddad, G. E., Blackwell, K. & Bikhazi, A. Regulation of insulin-like growth factor-1 by the renin-angiotensin system during regression of cardiac eccentric hypertrophy through angiotensin-converting enzyme inhibitor and AT1 antagonist. Can J Physiol Pharmacol 81, 142–149, doi:10.1139/y02-154 (2003).

49 Pickard, A. & McCance, D. J. IGF-Binding Protein 2 - Oncogene or Tumor Suppressor? Front Endocrinol (Lausanne) 6, 25, doi:10.3389/fendo.2015.00025 (2015).

50 Li, T. et al. IGFBP2: integrative hub of developmental and oncogenic signaling network. Oncogene 39, 2243–2257, doi:10.1038/s41388-020-1154-2 (2020).

51 Baron-Hay, S., Boyle, F., Ferrier, A. & Scott, C. Elevated serum insulin-like growth factor binding protein-2 as a prognostic marker in patients with ovarian cancer. Clin Cancer Res 10, 1796–1806, doi:10.1158/1078-0432.ccr-0672-2 (2004).

52 Beech, D. J., Parekh, N. & Pang, Y. Insulin-like growth factor-I receptor antagonism results in increased cytotoxicity of breast cancer cells to doxorubicin and taxol. Oncol Rep 8, 325–329, doi:10.3892/or.8.2.325 (2001).

53 Mamay, C. L., Mingo-Sion, A. M., Wolf, D. M., Molina, M. D. & Van Den Berg, C. L. An inhibitory function for JNK in the regulation of IGF-I signaling in breast cancer. Oncogene 22, 602–614, doi:10.1038/sj.onc.1206186 (2003).

54 Lu, Y., Zi, X., Zhao, Y., Mascarenhas, D. & Pollak, M. Insulin-like growth factor-I receptor signaling and resistance to trastuzumab (Herceptin). J Natl Cancer Inst 93, 1852–1857, doi:10.1093/jnci/93.24.1852 (2001).

55 Ajona, D. et al. Short-term starvation reduces IGF-1 levels to sensitize lung tumors to PD-1 immune checkpoint blockade. Nat Cancer 1, 75–85, doi:10.1038/s43018-019-0007-9 (2020).

