## supplemental materials for "Losartan rewires ovarian cancer tumor-immune microenvironment and suppresses IGF-1 to amplify chemo-immunotherapy sensitivity"

### Supplementary Materials and Methods:

**Patient characteristics.** All clinical samples were collected and utilized under approved Institutional Review Board protocols (#DFHCC07-049, 2014P002048, and 2022P000148) from patients who provided informed consent. The samples included either fresh frozen tumors and or paraffin blocks recovered from primary surgical cases from patients who were chemotherapy naïve with a confirmed diagnosis of high-grade serous ovarian cancer. Samples were either retrieved from the Vincent Center for Reproductive Biology - Massachusetts General Hospital (MGH) Gyn Repository or the MGH Department of Pathology archives.

**Cell lines and reagents.** BR5 and ID8 cells were cultured in Dulbecco's Mod. of Eagle's Medium (from Corning, Manassas, VA). The lentiviral vector carrying an expression cassette encoding Gaussia luciferase (Gluc) was obtained from the MGH vector core. BR5 and ID8 cells were transduced with Gluc at a multiplicity of infection (MOI) of 50 [1]. Recombinant angiotensin II was obtained from Enzo Life Sciences (Farmingdale, NY), and losartan was obtained from TCI America (Boston, MA). Anti-CD8 blocking antibody was obtained from BioXcell (clone 53-6.7, Lebanon, NH).

**Animal model.** All animal procedures followed the guidelines of the Public Health Service Policy on Humane Care of Laboratory Animals and approved by the Institutional Animal Care and Use Committee of the Massachusetts General Hospital. BR5-Gluc ($1x10^6$) or ID8-Gluc ($1x10^6$) tumor cells were injected *i.p.* into 8-12 weeks-old female FVB or C57/BL6 mice, respectively. Tumor growth was monitored by Gluc testing every 3 days until mice in the control group became moribund. All peritoneal tumors were excised and weighed and malignant ascites were aspirated and volume measured [2,3].

**Treatment protocols.**

*Losartan treatment.* Losartan (40 mg/kg) was administered by oral gavage once every day and continued until mice in the control group became moribund [4,5].

*Paclitaxel treatment.* Paclitaxel (obtained from MGH pharmacy, 5mg/kg) was administered *i.p.* in 100 µl of saline once per week until mice in the control group became moribund [2].

*Anti-PD1 treatment*. The anti-PD1 antibody or isotype control IgG (Bioxell, (200 µg/mice) was administrated *i.p.* every 3 days for a total of 4 dosages.

**Gaussia Luciferase measurement.** Tumor cell lines were infected with lentivirus encoding secreted Gaussia luciferase (Gluc), and the measurement of plasma Gluc was performed as previously described [6,7]. Briefly, 10 µl of blood was collected from the tail vein and mixed with 5 µl of 50 mM EDTA. Gluc activity was measured using a plate luminometer (MLX luminometer, Dynex Technologies, Chantilly, VA). The luminometer was set to automatically inject 100 µl of coelenterazine (CTZ, 100 mM, Nanolight, Pinetop, AZ) and photon counts were acquired for 10 sec. When the blood Gluc value reached $2x10^6$ RLU/s, mice were randomized into different treatment groups.

**mRNA extraction, quantitative RT-PCR, and RNASeq analysis.** The mRNA was extracted using the RNeasy kit (Qiagen, Redwood City, CA) according to the manufacturer's protocol. DNaseI digestion was added to the column during the extraction process as suggested by the manufacturer. RNA purity and quantity were estimated by NanoDrop ND-1000 Spectrophotometer 260/280. Quantitative RT-PCR (qRT-PCR) was done by SYBR®

Green-based, real-time cDNA or miRNAs PCR System (Qiagen). All qPCR and analysis were performed on a Stratagene MX 3000 qPCR System operating MXPro qPCR software (Stratagene, San Diego, CA). The data were analyzed with the web-based software package [2].

For bulk RNASeq analysis, tissues were homogenized using the Polytron PT1300 tissue homogenizer, followed by additional homogenization using a Qiashredder spin column. RNA from tumor tissues was extracted using the RNeasy Mini Kit (QIAGEN, Cambridge, MA). 1 μg of total RNA was sent to Molecular Biology Core Facilities, Dana-Farber Cancer Institute. RNASeq analysis was performed following the routine procedure in the Xu lab [8]. For bulk RNA-Seq data analysis, Fastqc software was used for raw data quality control. To remove adaptor contamination and low-quality bases we used Cutadapt. After data filtering, high-quality reads were mapped to the reference genome by using Hisat2, with each read at most 2 mismatches accepted. For human samples, we used the human reference genome hg19. For mice samples, we mapped the mice sample clean data to mouse reference genome mm9, as well as human reference genome hg19 by using Hisat2 with at most 2 mismatches for each read. After data mapping, we used Stringtie to assemble transcripts, as well as calculate the FPKM value for gene expression level. For each gene, we sum the expression value for all the transcripts. edgeR was used to identify differential expression genes. DAVID was used to do gene ontology and Kegg pathway enrichment annotation. GSEA was used to analyze the enriched gene sets. The DESeq2 package in R was used to determine the differentially expressed genes (DEGs) [9]. To control for False discovery rate (FDR) at 0.05, we used the Benjamini & Hochberg algorithm. ComplexHeatmap package was used to plot the heatmap [10]. The differentially expressed gene set was analyzed by Gene Set Enrichment Analysis software (GSEA, https://software.broadinstitute.org/ software/cprg/?q=node/14).

**Histology and immunohistochemistry.** For IHC and H&E-staining procedures, tumors were fixed in formalin and then embedded in paraffin. For IHC requiring frozen tissue, fresh tumor tissues were embedded in OCT compound (Miles, Inc., Elkhart, IN), frozen slowly on dry ice, and stored at -80°C. Tumor cell proliferation (PCNA, 1:800; Dako, Carpinteria, CA), apoptosis (TUNEL apoptosis detection kit; Millipore, Billerica, MA), and collagen I (Sigma-Aldrich, St. Louis, MO) were evaluated in paraffin sections. T cell infiltration was evaluated by staining for CD4 (1:200) and CD8 (1:50) T cells (both are from Cell Signaling, Waltham, MA).

*Image quantification:* For the CD4, CD8 T cells, PCNA, and TUNEL staining, positively stained cells are manually counted. For collagen I and anti-PD1 delivery staining, the positively stained area of all other histological staining was evaluated with digital quantitative image analysis using the open-source software ImageJ. Positive staining in 20 random fields/slides was quantified via automated built-in functions based on fluorescent pixel intensity after establishing a threshold to exclude background staining. Individual staining was quantified as area fractions of the tumor region of interest and reported as percentages [8].

**Protein expression analysis.**

*Western Blot.* Thirty micrograms of protein per sample were separated on 10% SDS-polyacrylamide gels [11]. Membranes were blotted with antibodies against total (1:1000) and phospho-Akt (1:200); total and phospho-S6 (1:1000 for both); total (1:1000) and phospho-ERK1/2 (1:1000). Antibodies were obtained from Cell Signaling (Danvers, MA). Membranes were blotted with beta-actin for equal loading control (1:5000, Sigma) [12].

*ELISA.* Plasma or protein extracted from snap-frozen tumors were diluted to 2 μg/μl concentration according to protein assay. Mouse IGF-1 cytokine levels were quantified using a Quantikine ELISA kit following the manufacturer's instructions (R&D Systems, Gaithersburg, MD). Inflammatory cytokine levels were quantified using multiplex enzyme-linked immunosorbent assay plates from Meso-Scale Discovery (MSD) following the manufacturer's instructions. Every sample was run in triplicate [13,14].

*Phospho-receptor kinase array.* The Mouse Phospho-Receptor Tyrosine Kinase (RTK) Array Kit (Catalog # ARY014, R&D Systems) was used to detect phosphorylation levels of 39 RTKs following the manufacturer's protocol. Two hundred micrograms of total protein lysate from each sample were used per membrane. Protein concentration was determined using Pierce BCA Protein Assay Reagent (ThermoFisher Scientific). Following processing, each array membrane was exposed to X-ray film for 1 h and developed using a JPI X-ray film processor. The relative density of positive array hits was analyzed using ImageJ v1.53e software gel analysis and background subtraction was applied using the included negative PBS control.

**Flow cytometry**. To characterize the tumor-infiltrating immune cells, as well as profile immune cells in the ascites, the peritoneal tumors were removed and homogenized in Roswell Park Memorial Institute (RPMI) medium, filtered through a 40-μm nylon cell strainer (BD Falcon). Tumor cells and ascites were centrifuged at 1000 rpm, and resuspended in 4 mL 30% Percoll. The cells were overlaid onto a Percoll gradient (30%/37%/60%) and centrifuged at 1200 rpm for 20 minutes. Lymphocytes were collected from the 37%/60% interface and washed twice in phosphate-buffered saline (PBS), then stimulated for 6 hours with RPMI + phorbol myristate acetate (PMA) + ionomycin (Sigma-Aldrich) + Golgi-stop (BD Biosciences) at 37°C. After incubation, the cells were washed with PBS and stained with anti-CD45 (clone 30-F11), anti-CD4 (clone RM4-5), anti-CD8 (clone 53-6.7), anti-Foxp3 for Treg (clone. D6O8R), anti-NK1.1 (clone PK136), anti-Gr1 for MDSC (clone RB6-8C5), anti-CD11b (clone m1/70) and anti-F4/80 (clone BM8) for Macrophage, anti-iNOS for M1 (clone CXNFT) and anti-Arginase I for M2 (clone D4E3M) anti-CD86 for activated dendritic cells (clone GL-1), and anti- interferon (IFN-γ) (clone XMG1.2), anti-TNFα (clone MP6-XT22) , anti-Granzyme B (clone GB11). Appropriate isotype controls were used. The cells were analyzed by fluorescence-activated cell sorting (FACS) analysis using a FACSCalibur flow cytometer (BD Biosciences). All antibodies were purchased from eBioscience. Data were analyzed with FlowJo software.

**Plasmid constructs.** Murine *Igf-1* (GenBank: NM_010512) was amplified from murine liver cDNA library using primers: Forward: 5'- GGA TCC AAG CTT ATG GGG AAA ATC AGC AGC CT -3'; Reverse: 5'- GAA TTC GCG GCC GCC TAC TTG TGT TCT TCA AAT G -3'. The amplified fragment was cloned into pBEAK33 vector (from Brian Seed laboratory), and its sequence was confirmed. After transfection of the plasmid, transfected cells were pooled to avoid clonal variation and screened for their overexpression of IGF-1 by qRT-PCR and ELISA (R&D Systems).

**Isolation of TAMs and peritoneal macrophages**. Tumor-associated macrophages were isolated by fluorescence-activated cell sorting. Briefly, tumor tissues were mechanically chopped and enzymatically digested for 1 hour at 37°C with 3mg/ml collagenase A (Roche) in a serum-free DMEM medium. Digestion was stopped by the addition of DMEM supplemented with 8% FBS and the suspension was disaggregated through a 70 μm cell strainer. Single-cell suspension was stained for 20 min at 4°C with anti-mouse F4/80 (1:200, BM8; eBioscience). Sorting of F4/80$^+$ cells was performed on a BD FACS sorter. Peritoneal macrophages were collected by peritoneal lavage from mice given an intraperitoneal injection of 2 ml of thioglycolate broth (Sigma) following our previously published protocol [15]. Briefly, 4 days after thioglycollate injection, peritoneal macrophages were harvested and washed with $Ca^{2+}$- and $Mg^{2+}$-free PBS, and suspended in 10% DMEM for culture and in vitro treatment [15].

**MTT assay.** Five thousand cells were seeded into 38 mm$^2$ wells of flat-bottomed 96-well plates in triplicate and allowed to adhere overnight. 3-(4,5-Dimethylthiazol-2-yl)-2,5-diphenyltetrazolium bromide (MTT, 5 mg/mL, Sigma Chemical) was prepared in PBS. The number of metabolically active cells was determined by MTT assay [16].

**Statistical analyses.** Significant differences between the two groups were analyzed using the Student's t-test (two-tailed). The incidence of ascites was analyzed using the two-sided Fisher's exact test. Histological analysis of gene expression in patient OvCa tumor tissues was statistically analyzed using statistics software (Statistical Package for the Social Sciences Statistics, Version 20.0; IBM).

**Supplementary Figures**

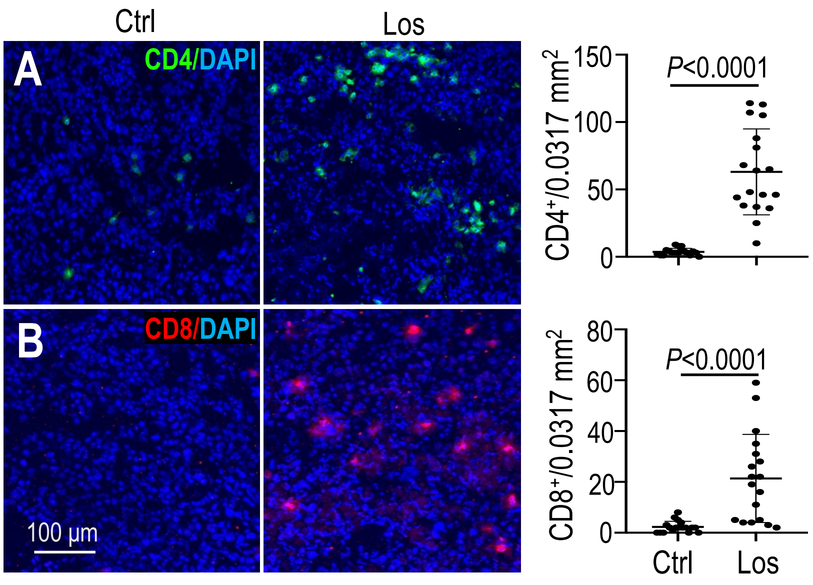


**Figure S1. Histological confirmation of losartan-induced tumor-infiltrating CD4 and CD8 T cells.** In the BR5 model, peritoneal tumors collected on day 35 post-implantation were immunofluorescently stained **(A)** CD4, and **(B)** CD8. The number of positively stained cells/0.0317 $mm^2$ area was manually counted in 20 random fields. Data presented as Mean ± SD.

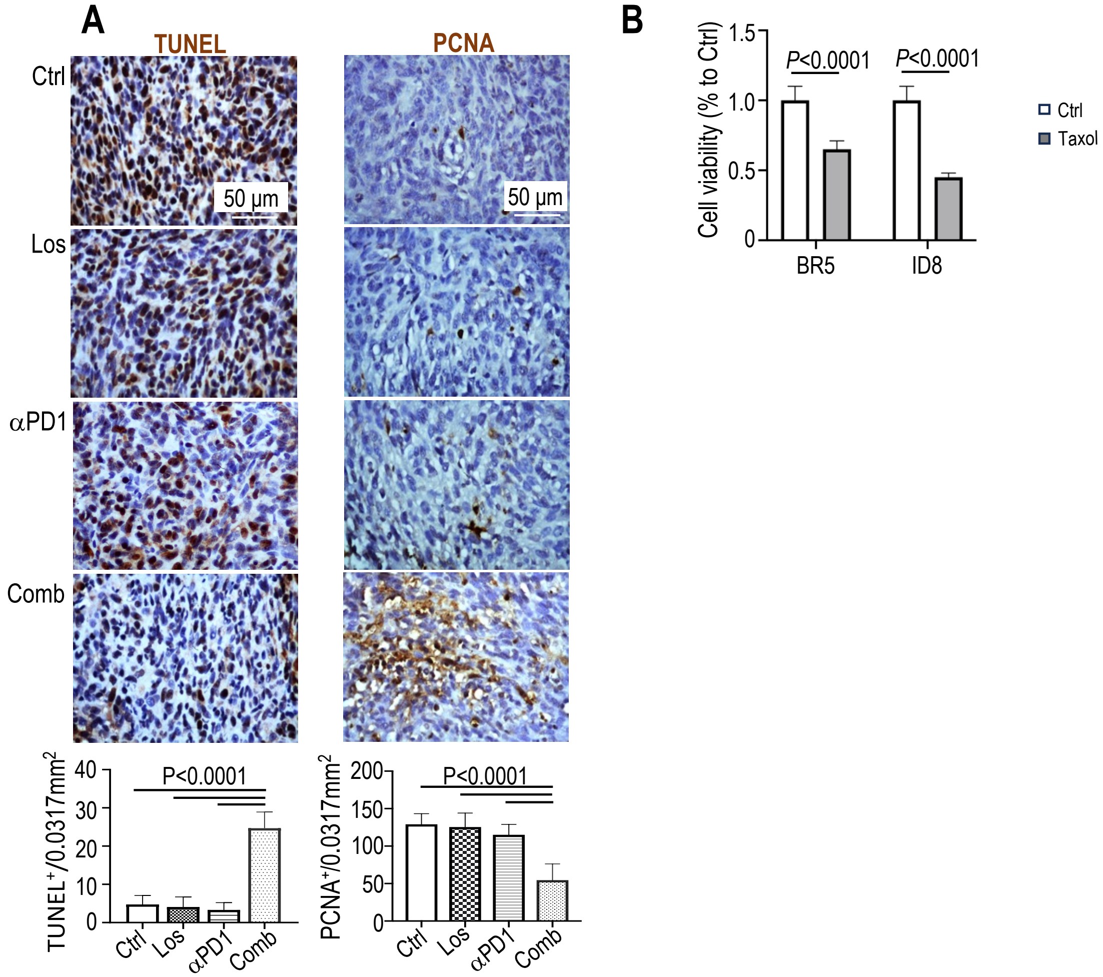


**Figure S2. Combined losartan treatment improves immunotherapy efficacy in OvCa models**. **(A)** Peritoneal BR5 tumors were collected from mice in the treatment groups receiving: i) control (saline), ii) losartan (40 mg/kg, QD), iii) αPD1 (200 μg for 4 treatments), v) losartan+αPD1. Tumor cell apoptosis was evaluated by TUNEL staining and tumor cell proliferation was evaluated using PCNA IHC staining. The number of TUNEL and PCNA positively stained cells was manually counted in 20 random fields (0.0317 $mm^2$/ea). Data presented as Mean ± SD. N=3 mice/group. **(B)** BR5 and ID8 cells were treated with saline (control, Ctrl) or Taxol (10 nM) Taxol for 3 days, cell viability was evaluated by MTT assay. Data presented as Mean ± SD.

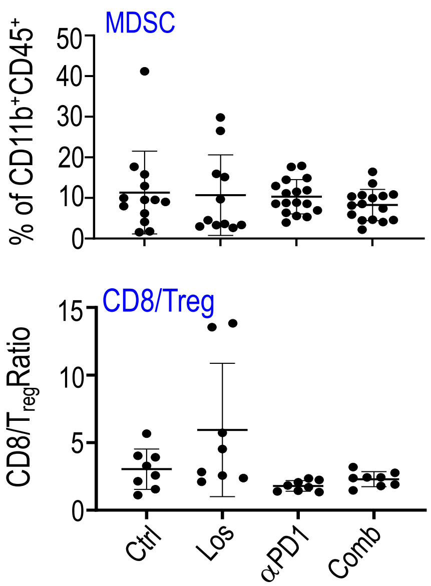


**Figure S3. Effect of losartan treatment on immune cell infiltration in mouse OvCa models.** Peritoneal BR5 tumors were collected from mice in the treatment groups receiving: i) control (saline), ii) losartan (40 mg/kg, QD), iii) αPD1 (200 μg for 4 treatments), and v) losartan+αPD1. Flow cytometry analysis of MDSC and the ratio of CD8/Treg. Data presented are mean ± SEM.

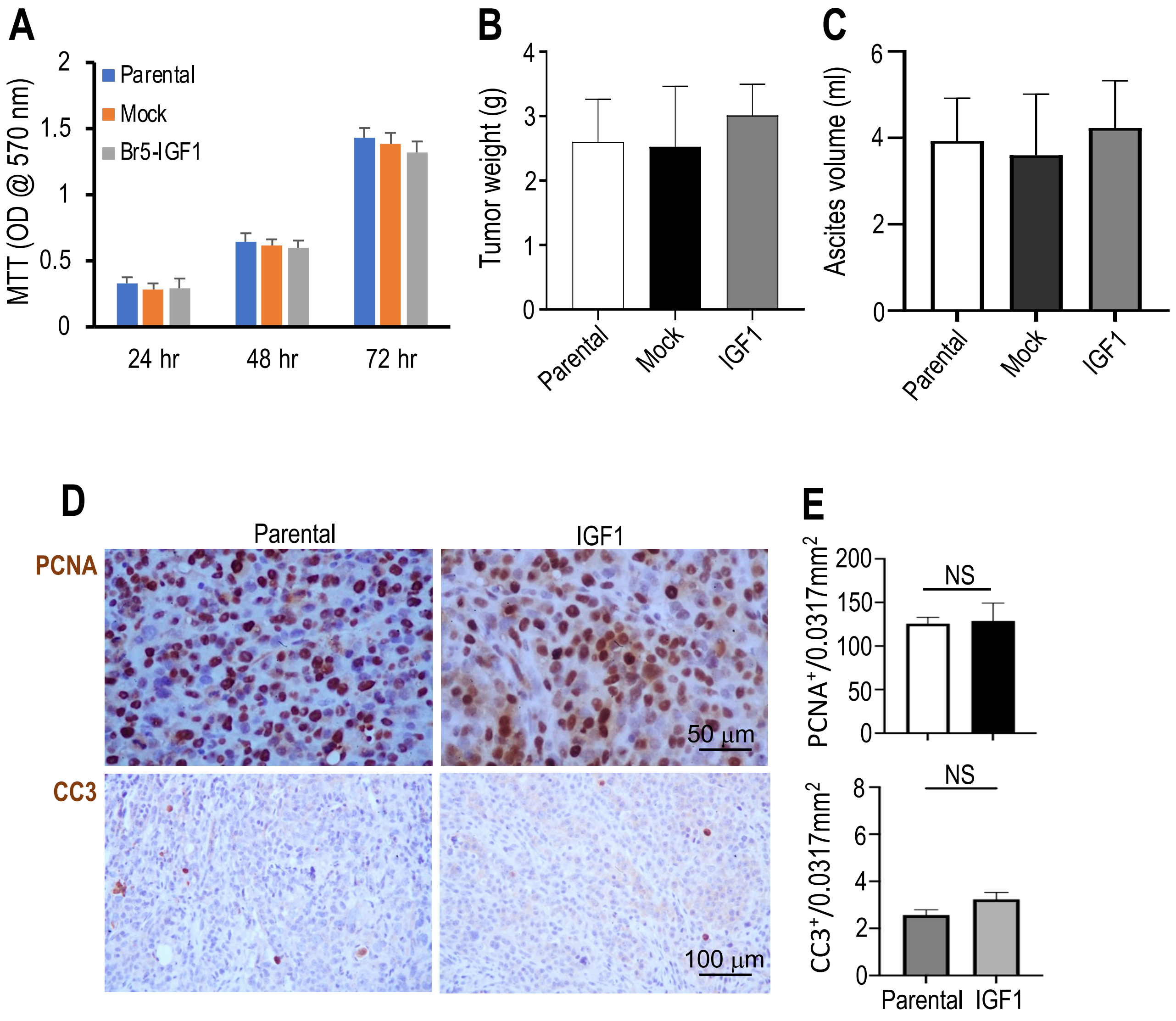


**Figure S4. IGF-1 overexpression does not affect OvCa tumor growth. (A)** Parental, mock-transfected, and IGF-1 overexpressing BR5 cells were plated in a 96-well plate and an MTT assay was performed to evaluate cell viability at 24, 48, and 72 hours. **(B-C)** Parental, mock-transfected, and IGF-1 overexpressing BR5 cells were implanted into the peritoneal cavity of FVB mice. When mice became moribund, **(B)** peritoneal tumor weight and **(C)** ascites volume were measured. N=6/ea, Data presented are mean ± SEM. **(D-E)** Tumor cell proliferation was evaluated using PCNA IHC staining and tumor cell apoptosis was evaluated by cleaved caspase-3 (CC3) staining. The number of PCNA and CC3 positively stained cells was manually counted in 20 random fields (0.0317 $mm^2$/ea). Data presented as Mean ± SD. N=3 tumors/group.

**REFERENCES CITED:**


1 Zhao, Y. *et al.* Losartan treatment enhances chemotherapy efficacy and reduces ascites in ovarian cancer models by normalizing the tumor stroma. *Proc Natl Acad Sci U S A*, pii:201818357. 201818310.201811073/pnas.1818357116, doi:10.1073/pnas.1818357116 (2019).

2 Liao, S. *et al.* TGF-beta blockade controls ascites by preventing abnormalization of lymphatic vessels in orthotopic human ovarian carcinoma models. *Clin Cancer Res* **17**, 1415-1424, doi:10.1158/1078-0432.CCR-10-2429 (2011).

3 Xu, L. & Fidler, I. J. Acidic pH-induced elevation in interleukin 8 expression by human ovarian carcinoma cells. *Cancer Res* **60**, 4610-4616 (2000).

4 Chauhan, V. P. *et al.* Angiotensin inhibition enhances drug delivery and potentiates chemotherapy by decompressing tumour blood vessels. *Nat Commun* **4**, 2516-2527, doi:10.1038/ncomms3516 (2013).

5 Diop-Frimpong, B., Chauhan, V. P., Krane, S., Boucher, Y. & Jain, R. K. Losartan inhibits collagen I synthesis and improves the distribution and efficacy of nanotherapeutics in tumors. *Proc Natl Acad Sci U S A* **108**, 2909-2914, doi:10.1073/pnas.1018892108 (2011).

6 Gao, X. *et al.* Anti-VEGF treatment improves neurological function and augments radiation response in NF2 schwannoma model. *Proc Natl Acad Sci U S A* **112**, 14676-14681, doi:10.1073/pnas.1512570112 (2015).

7 Zhao, Y. *et al.* Targeting the cMET pathway augments radiation response without adverse effect on hearing in NF2 schwannoma models. *Proc Natl Acad Sci U S A* **115**, E2077-E2084, doi:10.1073/pnas.1719966115 (2018).

8 Wu, L. *et al.* Losartan prevents tumor-induced hearing loss and augments radiation efficacy in NF2 schwannoma rodent models. *Sci Transl Med* **13**, doi:10.1126/scitranslmed.abd4816 (2021).

9 Love, M. I., Huber, W. & Anders, S. Moderated estimation of fold change and dispersion for RNA-seq data with DESeq2. *Genome Biol* **15**, 550, doi:10.1186/s13059-014-0550-8 (2014).

10 Gu, Z., Eils, R. & Schlesner, M. Complex heatmaps reveal patterns and correlations in multidimensional genomic data. *Bioinformatics* **32**, 2847-2849, doi:10.1093/bioinformatics/btw313 (2016).

11 Zhao, Y. *et al.* Losartan treatment enhances chemotherapy efficacy and reduces ascites in ovarian cancer models by normalizing the tumor stroma. *Proc Natl Acad Sci U S A* **116**, 2210-2219, doi:10.1073/pnas.1818357116 (2019).

12 Xu, L., Pathak, P. S. & Fukumura, D. Hypoxia-induced activation of p38 mitogen-activated protein kinase and phosphatidylinositol 3'-kinase signaling pathways contributes to expression of interleukin 8 in human ovarian carcinoma cells. *Clin Cancer Res* **10**, 701-707, doi:10.1158/1078-0432.ccr-0953-03 (2004).

13 Xu, L. *et al.* Direct evidence that bevacizumab, an anti-VEGF antibody, up-regulates SDF1alpha, CXCR4, CXCL6, and neuropilin 1 in tumors from patients with rectal cancer. *Cancer Res* **69**, 7905-7910, doi:10.1158/0008-5472.CAN-09-2099 (2009).

14 Xu, L. *et al.* Placenta growth factor overexpression inhibits tumor growth, angiogenesis, and metastasis by depleting vascular endothelial growth factor homodimers in orthotopic mouse models. *Cancer Res* **66**, 3971-3977, doi:10.1158/0008-5472.CAN-04-3085 (2006).

15 Xu, L., Xie, K. & Fidler, I. J. Therapy of human ovarian cancer by transfection with the murine interferon beta gene: role of macrophage-inducible nitric oxide synthase. *Hum Gene Ther* **9**, 2699-2708, doi:10.1089/hum.1998.9.18-2699 (1998).

16 Xu, L., Tong, R., Cochran, D. M. & Jain, R. K. Blocking platelet-derived growth factor-D/platelet-derived growth factor receptor beta signaling inhibits human renal cell carcinoma progression in an orthotopic mouse model. *Cancer research* **65**, 5711-5719, doi:10.1158/0008-5472.CAN-04-4313 (2005).
